# Comparative systems analysis of the secretome of the opportunistic pathogen *Aspergillus fumigatus* and other *Aspergillus* species

**DOI:** 10.1101/230953

**Authors:** R.P. Vivek-Ananth, Karthikeyan Mohanraj, Muralidharan Vandanashree, Anupam Jhingran, James P. Craig, Areejit Samal

**Affiliations:** The Institute of Mathematical Sciences, Homi Bhabha National Institute, Chennai 600113, India; Stony Brook University, Stony Brook, New York 11794-3369, USA

## Abstract

*Aspergillus fumigatus* and multiple other *Aspergillus* species cause a wide range of lung infections, collectively termed aspergillosis. *Aspergilli* are ubiquitous in environment with healthy immune systems routinely eliminating inhaled conidia, however, *Aspergilli* can become an opportunistic pathogen in immune-compromised patients. The aspergillosis mortality rate and emergence of drug-resistance reveals an urgent need to identify novel targets. Secreted and cell membrane proteins play a critical role in fungal-host interactions and pathogenesis. Using a computational pipeline integrating data from high-throughput experiments and bioinformatic predictions, we have identified secreted and cell membrane proteins in ten *Aspergillus* species known to cause aspergillosis. Small secreted and effector-like proteins similar to agents of fungal-plant pathogenesis were also identified within each secretome. A comparison with humans revealed that at least 70% of *Aspergillus* secretomes has no sequence similarity with the human proteome. An analysis of antigenic qualities of *Aspergillus* proteins revealed that the secretome is significantly more antigenic than cell membrane proteins or the complete proteome. Finally, overlaying an expression dataset, four *A. fumigatus* proteins upregulated during infection and with available structures, were found to be structurally similar to known drug target proteins in other organisms, and were able to dock *in silico* with the respective drug.

## Introduction

Aspergillosis is an umbrella term for a wide array of infections caused by multiple *Aspergillus* species. The majority of reported aspergillosis cases originate from ten species: *Aspergillus fumigatus, Aspergillus flavus, Aspergillus niger, Aspergillus terreus, Aspergillus versicolor, Aspergillus lentulus, Aspergillus nidulans, Aspergillus glaucus, Aspergillus oryzae,* and *Aspergillus ustus*^1-3^. *A. fumigatus* alone is responsible for over 90% of the reported cases, followed by *A. flavus, A. niger, A. terreus* and *A. versicolor*^2,4,5^ The disease is an increasing concern for immune-compromised individuals, putting at risk patients with neutropenia, allogenic stem cell transplantation, organ transplantation, and acquired immunodeficiency syndrome (AIDS), among others^6^. High-risk patients can see mortality rates between 40-90%^7^. Compounding the issue is the limited number of drugs available to treat aspergillosis, and their widespread use has led to the emergence of drug-resistant strains, creating the pressing need for novel drugs, biomarkers and preventive therapies such as vaccines^8^.

Aspergillosis is caused by an opportunistic pathogen. Under normal conditions, *Aspergilli* are saprotrophic fungi which are present in soil and decaying organic matter, thereby, playing a fundamental role in nitrogen and carbon recycling. They have a broad geographical range with colonies typically spread through microscopic airborne conidia. As *Aspergillus* conidia are ubiquitous in the atmosphere, humans often inhale several hundred spores daily. While these spores are quickly eliminated by a healthy immune system^9^, in an immune-compromised host, the fungus becomes an opportunistic pathogen and overwhelms the weakened defences. The underlying mechanisms behind the successful initiation of pathogenesis by *Aspergilli* remain unclear. However, studies have shown that in immune-compromised hosts, *A. fumigatus* can reach the respiratory epithelia upon inhalation where it secretes proteases and secondary metabolites, specifically gliotoxin, which aids in colonizing healthy lung tissue^10,11^. Proteases secreted by *A. fumigatus* have also been implicated in establishing infection by morphologically altering respiratory epithelial cells^11,12^. In addition, *A. fumigatus* secretes a diverse array of catabolic enzymes that enable degradation of macromolecular biopolymers like elastin and collagen, which are present in large quantities in lung tissue, for uptake of nutrients^13^. Apart from secreted proteins, cell membrane and cell wall proteins also play a crucial role in interactions with the host immune system and establishing infection^10^. Thus, the diverse set of secreted and cell membrane proteins of *A. fumigatus* and other *Aspergillus* species play a vital role in pathogenesis.

Given the central role played by the secretome and cell membrane proteins of *Aspergillus* species during human pathogenesis, they are potential candidates for identifying new biomarkers, vaccines and druggable targets. This is especially urgent as prophylactic use of antifungal therapies has led to the emergence of resistance in many *Aspergillus* species to antifungal drugs^8,14,15^. Prospecting the secretome and cell membrane proteins of *Aspergillus* species for druggable targets is advantageous, since any identified target will be exposed to the extracellular space. In addition, targeting extracellular rather than intracellular proteins will provide fewer mechanisms for resistance to develop in the pathogen.

This study aims to identify and analyze the secretome and cell membrane proteins of *A. fumigatus* and nine other *Aspergillus* species known to cause aspergillosis, namely, *A. flavus, A. niger, A. terreus, A. versicolor, A. lentulus, A. nidulans, A. glaucus, A. oryzae,* and *A. ustus.* To our knowledge, this is the first dedicated effort to characterize the secretome and cell membrane proteins of *Aspergillus* species with an aim to understand their role in pathogenesis and to analyze them as potential druggable proteins. While previous studies such as FSD^16^, FunSecKB2^17^ and SECRETOOL^18^ have developed computational pipelines for fungal secretome prediction, they rely solely on bioinformatic predictions even though many experimental high-throughput proteomic datasets in *Aspergillus* species are available. Thus, we have designed a computational pipeline which systematically integrates experimental high-throughput proteomic datasets, UniProt^19^ annotations with experimental evidence, and predictions from bioinformatic tools to identify a comprehensive set of secreted extracellular proteins and cell membrane proteins in *Aspergillus* species. Furthermore, small secreted and effector-like proteins^20-23^ were identified in *Aspergillus* species, and the set of secreted and cell membrane proteins were analyzed for the abundance of antigenic regions (AAR)^24^ and similarity to known drug target proteins from DrugBank^25^. Finally, analysis of a published gene expression dataset^26^ for *A. fumigatus,* led to identification of secreted and cell membrane proteins which are upregulated under pathogenesis and have no human homologs, and such candidates were taken for further druggability analysis by computational docking experiments, thereby identifying pre-existing drugs used against other pathogens as candidates for repurposing against aspergillosis.

## Results and discussion

### Identification of the secretome from high-throughput experimental studies

An extensive literature search was undertaken to compile high-throughput experimental studies that have characterized the secretome of *Aspergillus* species. A systematic search led to a comprehensive list of 46 high-throughput proteomic studies^27-73^ on the secretome of 6 *Aspergillus* species analyzed here. Note that many high-throughput studies report proteins with transmembrane (TM) domains as part of secretome but a few studies have specifically identified cell membrane and cell wall associated proteins along with the secretome^30,36^. Our compilation of high-throughput proteomic studies led to experimentally identified lists of 437, 364, 454, 96, 469 and 171 secreted proteins in *A. fumigatus^27-41^,* A.favus^30,42-45^, *A. niger*^30,46-58^, *A. terreus^30,59,60^, A. nidulans*^58,61-65^ and *A. oryzae*^57,66-73^, respectively (Methods; Supplementary Table S1).

### Computational pipeline for fungal secretome prediction

High-throughput experimental studies that identify secretome can be limited by the detection technology used or the small set of experimental conditions assessed. Such secretomic studies mainly focus on identifying the proteins secreted to the extracellular matrix and not on proteins incorporated into the cell membrane by the eukaryotic secretion pathway. Since cell membrane proteins, including integral proteins, are localized on the cell surface, they can serve as excellent targets for drugs or vaccines. Here we have designed a computational pipeline to predict both secreted extracellular and cell membrane proteins in fungi. Importantly, our pipeline also incorporates and prioritizes available information on the experimentally identified secreted and cell membrane proteins. Furthermore, our pipeline subdivides the secretome into two subsets, proteins that follow the classical secretion pathway through the endoplasmic reticulum (ER) and those that exit the cell boundary through a non-classical secretion pathway. Figure 1 contains a flowchart of our prediction pipeline.

**Figure 1:**
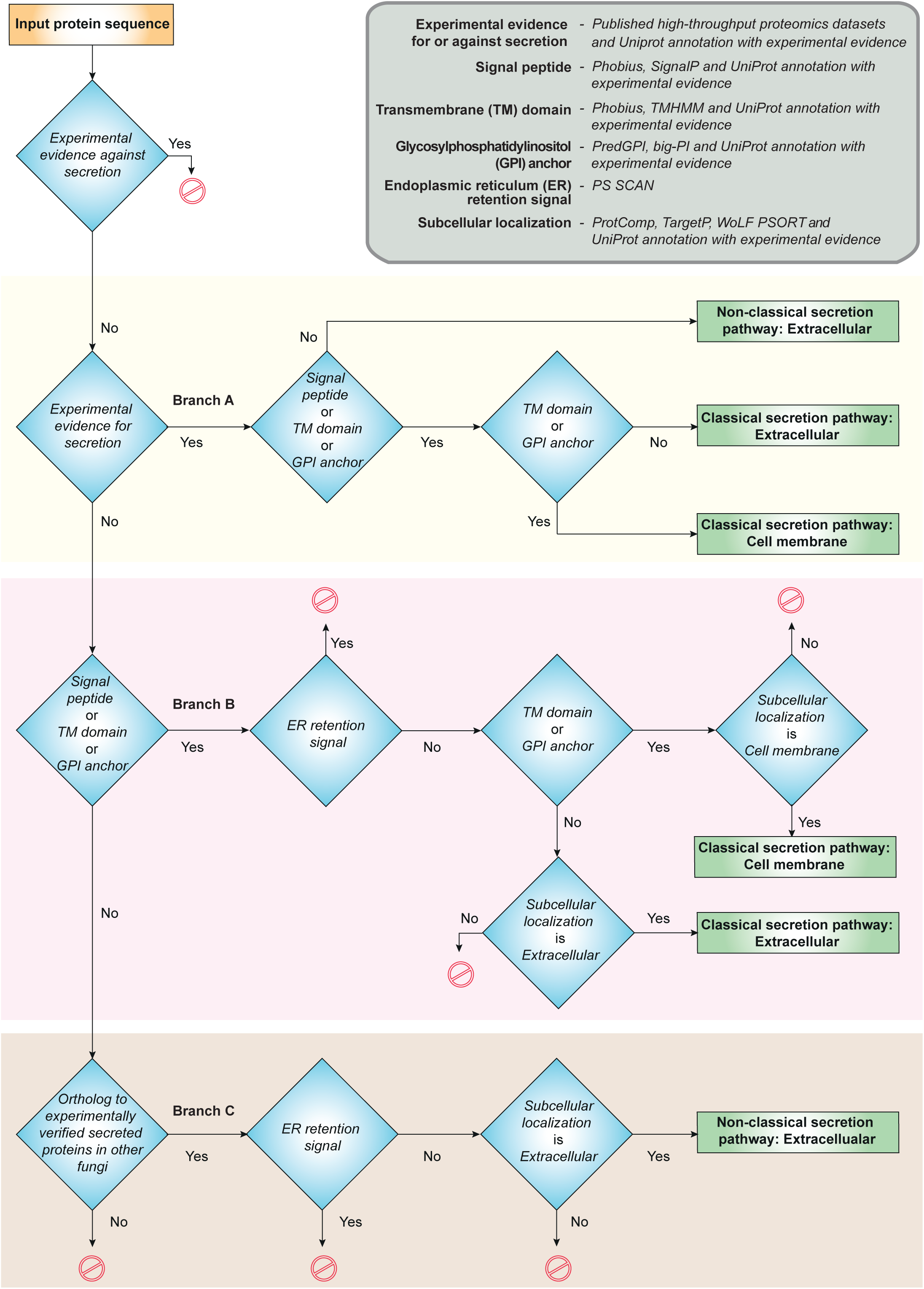
Flowchart of the computational prediction pipeline for identifying secreted extracellular proteins (secretome) and cell membrane proteins in *Aspergillus* species. Note that this prediction pipeline can be employed to identify secretome and cell membrane proteins in any fungi with sequenced genome.

Our pipeline starts from the complete proteome of a fungus. Initially, intracellular proteins based on UniProt^19^ annotation with experimental evidence were removed from later analysis. Subsequently, the remaining proteins without experimental evidence for intracellular localization were classified into two mutually exclusive categories. The first category contained secreted or cell membrane proteins with experimental evidence from compiled list of high-throughput proteomic studies or UniProt, and the second category contained proteins without experimental evidence of secretion to the extracellular matrix or localization to the cell membrane.

The first category of proteins with experimental evidence were assessed for signal peptide, Glycosylphosphatidylinositol (GPI) anchor or TM domain, confirming passage through the classical pathway (Branch A in Figure 1; Methods). Next, the presence of GPI anchor or TM domain in proteins sorted by classical pathway is used to separate the cell membrane from extracellular proteins. Next, the proteins without a signal peptide, GPI anchor and TM domain but with experimental evidence of secretion to extracellular matrix were assigned to non-classical pathway (Branch A in Figure 1).

The second category of proteins without experimental evidence were screened based on predictions from computational tools as follows. Firstly, the second category of proteins were screened for signal peptide, GPI anchor or TM domain, suggesting translocation into the ER and passage through classical pathway. Next, the proteins predicted to have a signal peptide, GPI anchor or TM domain but also with an ER retention signal were removed from later analysis (Branch B in Figure 1; Methods). Next, the proteins predicted to have GPI anchor or TM domain along with subcellular localization as cell membrane were classified as cell membrane proteins, and proteins without GPI anchor and TM domain along with subcellular localization as extracellular were classified as extracellular proteins sorted by classical pathway (Branch B in Figure 1).

Lastly, the subset of the second category of proteins which lack signal peptide, GPI anchor and TM domain, were assessed for presence of orthologs in the list of secreted proteins with experimental evidence in other fungi (Methods; Branch C in Figure 1). Next, those proteins in the subset which are orthologs of experimentally identified secreted proteins in other fungi were assessed for an ER retention signal and their predicted subcellular localization, and those without an ER retention signal and predicted subcellular localization as extracellular were classified as extracellular proteins secreted through a non-classical secretion pathway (Branch C in Figure 1). To our knowledge, SecretomeP^74,75^ is the only prediction tool for non-classical secretion pathway, however, SecretomeP^74,75^ is designed for bacteria and mammals and not for fungi. Still FSD^16^ has employed SecretomeP^74,75^ to predict approximately 40% of the proteome in several fungi as secreted via a non-classical secretion pathway which clearly is an overestimation. For example, FSD predicts 3546 *A. fumigatus* proteins, which is 35% of the proteome, to be secreted by a non-classical pathway. Thus, we employ here an alternate ortholog-based method to predict proteins secreted via a non-classical pathway.

Using this pipeline, we identified comprehensive sets of secreted and cell membrane proteins in ten *Aspergillus* species causing aspergillosis; Figure 2 displays the size of each set, Supplementary Table S2 lists the proteins in each set, and Supplementary Table S3 gives detailed annotations such as protein family, conserved domain, carbohydrate-binding modules and gene ontology (GO) terms for proteins in each set. Additionally, our pipeline provides a refined view of the protein sorting mechanisms in *Aspergillus* species by classifying secreted proteins into classical and non-classical pathway. In *A. fumigatus,* the predicted set of secreted and cell membrane proteins contained 662 and 1129 proteins, respectively, representing 6.7% and 11.5% of the proteome. Among the 662 proteins in *A. fumigatus* secretome, 64 were predicted to be secreted by a non-classical pathway. Figures 3 shows the significantly enriched GO biological processes in the predicted secretome and cell membrane proteins of *A. fumigatus.* Within the *A. fumigatus* secretome, 598 proteins sorted by classical pathway have GO annotations related to carbohydrate metabolism, proteolysis and cell wall modification, and 64 proteins secreted by a non-classical pathway have GO annotations related to carbohydrate metabolism, response to reactive oxygen species (ROS), alcohol degradation and gliotoxin metabolism. Many of the GO processes annotated with *A. fumigatus* proteins secreted by non-classical pathway are associated with virulence. Especially, gliotoxin has immunosuppressive properties and is suspected to be an important virulence factor in *Aspergillus* pathogenesis^76-79^. This suggests that non-classical pathways may be important for the secretion of virulence factors in *A. fumigatus*.

**Figure 2:**
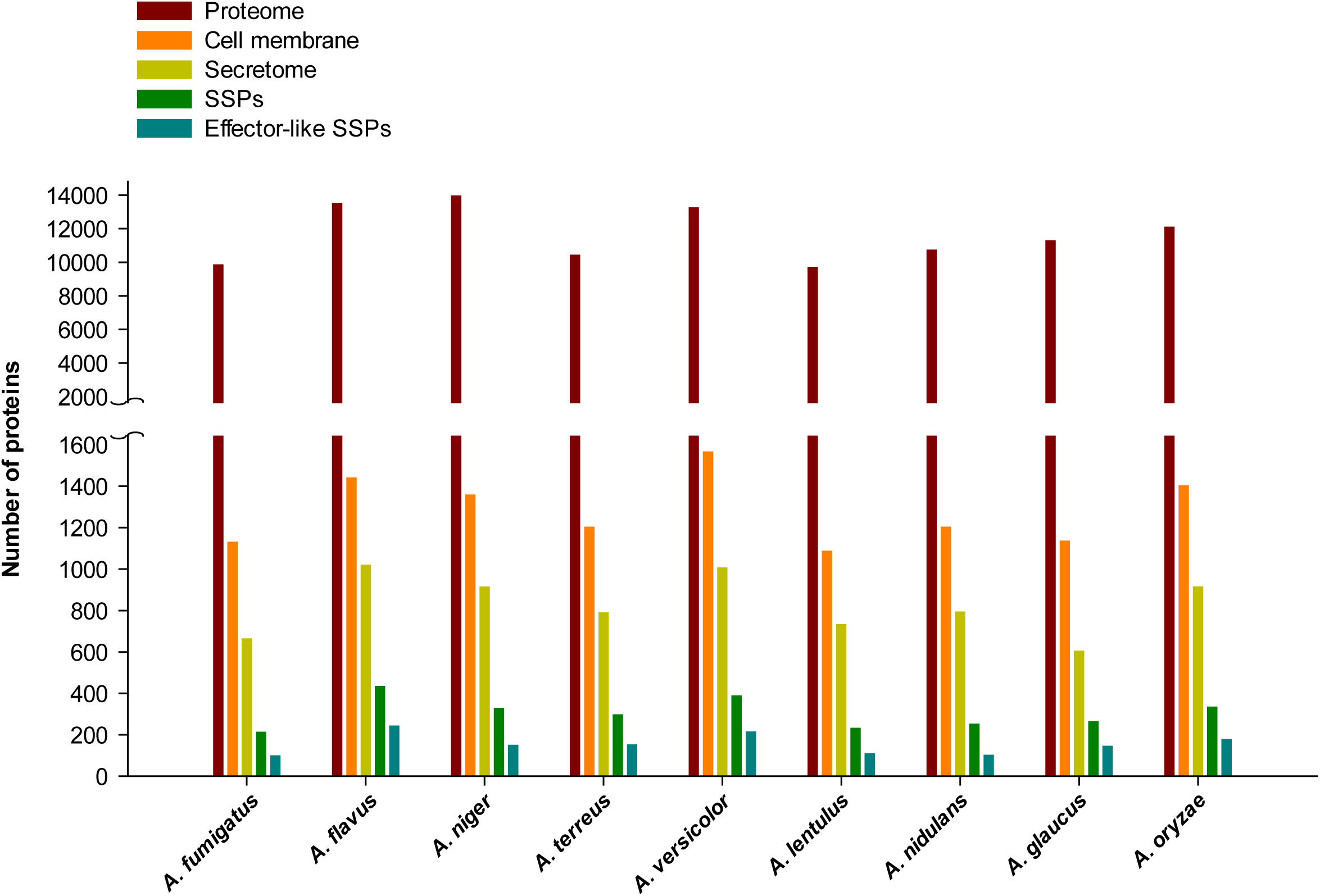
Number of proteins in the complete proteome, set of cell membrane proteins, set of secreted extracellular proteins, set of small secreted proteins (SSPs) and set of effector-like SSPs identified by our computational pipeline in *Aspergillus* species considered here.

**Figure 3:**
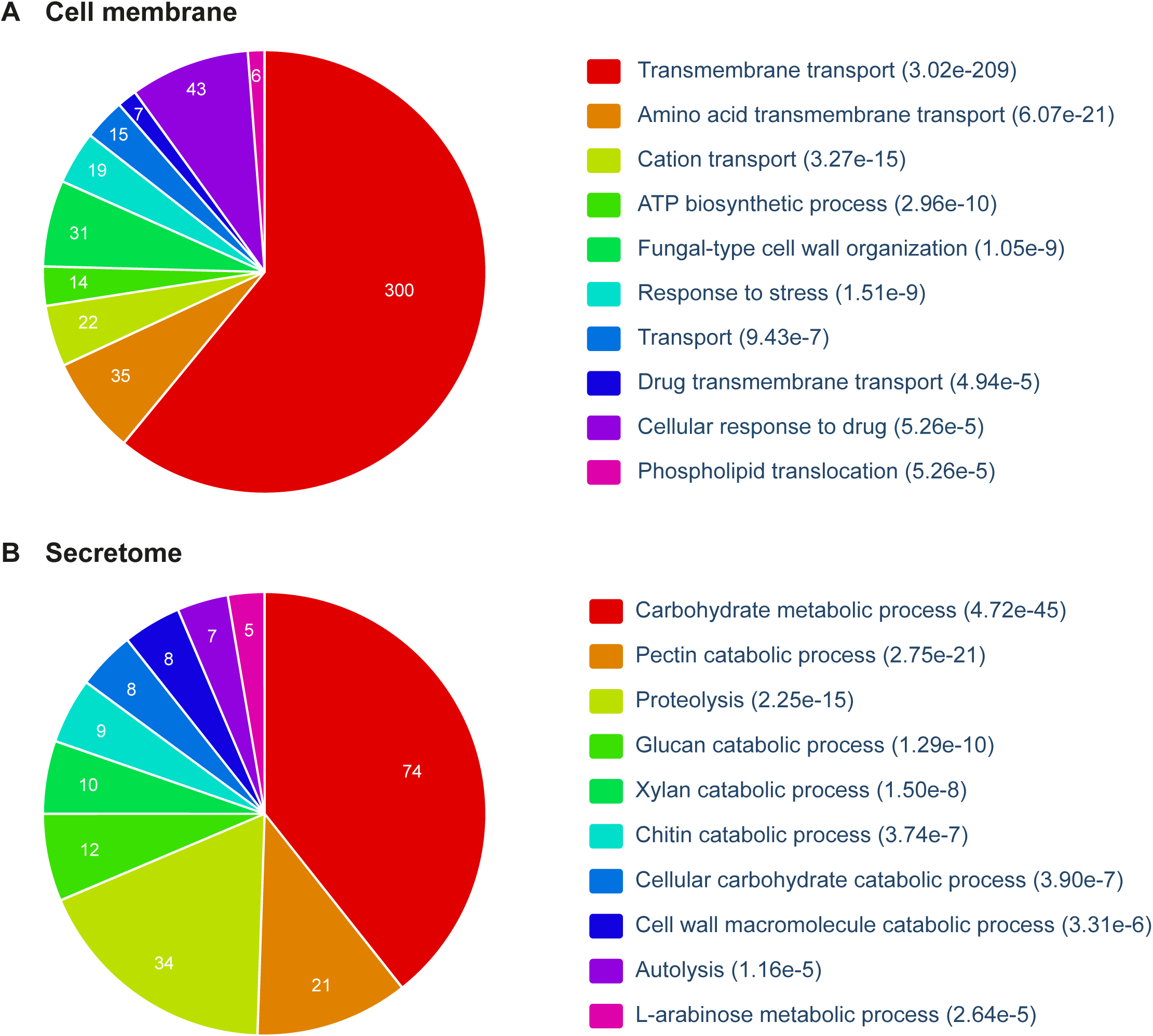
Gene ontology (GO) enrichment analysis for **(a)** cell membrane proteins and **(b)** secreted extracellular proteins in *A. fumigatus* Af293. In each case, the top 10 significantly enriched biological processes are shown along with p-value computed using Benjamini-Hochberg procedure. The GO enrichment analysis was performed using FungiFun2^130^ webserver (Methods).

### Small secreted and effector-like proteins

Within the fungal secretome, small secreted proteins (SSPs) with sequence length less than 300 amino acids have been widely studied for their role in fungal-plant pathogenesis^20,21^. A few of the SSPs have been found to act as effectors that play a central role in establishing plant infection^22,23^. Typically, fungal effector proteins do not share conserved domains which renders effector prediction a challenge^80,81^. Still, fungal effectors often share certain sequence characteristics such as size, cysteine content, or small motifs identified in fungi and oomycetes^80,81^. A recent study^82^ compared the SSPs across eight *Aspergillus* species focusing on plant biomass degradation. However, the role of SSPs and effector-like proteins in fungal-human pathogenesis, including aspergillosis, remains largely unanswered^83^. Thus, we have identified SSPs and effector-like proteins within the secretomes of *Aspergillus* species to enable discovery of potential virulence proteins (Supplementary Table S4). Note that effector-like proteins of *Aspergillus* species were predicted based on EffectorP^81^ predictions or cysteine content of SSPs (Methods). We found that SSPs and effector-like proteins account for more than 30% and 10%, respectively, of the secretomes in *Aspergillus* species (Figure 2). Specifically, 96 effector-like proteins were predicted among 210 SSPs in *A. fumigatus* secretome, of which, 44 do not have conserved Pfam domains and 4 contain one of the known fungal or oomycete effector motifs^80^, making them intriguing candidates for future virulence experiments.

### Upregulated secretome and cell membrane proteins in *A. fumigatus* during pathogenesis

To place the identified secretome and cell membrane proteins of *A. fumigatus* within the context of aspergillosis, we overlaid a previously published gene expression dataset^26^. Significantly differentially expressed and upregulated genes coding for secreted and cell membrane proteins may suggest a vital path or key towards pathogenesis. Note that several gene expression studies in *A. fumigatus* have been published on various cell cultures. However, due to the difficulty in obtaining sufficient high-quality RNA directly from sites of infection, gene expression studies of *A. fumigatus* in live animal models are limited. Thus, we selected for our analysis a published microarray dataset^26^ from a murine lung model which provided one of the first transcriptional snapshots of *A. fumigatus* during initiation of mammalian infection.

In this dataset^26^, *A. fumigatus* genes significantly upregulated over 2-fold during pathogenesis when compared to control conditions were selected for further analysis, totaling, 1264 proteins (12.9%) of the proteome. Within the predicted *A. fumigatus* secretome, 121 out of 662 proteins (18.3%) were upregulated over 2-fold during pathogenesis, indicating that a larger fraction of secretome is employed during pathogenesis. Moreover, 113 out of the 121 upregulated proteins in the *A. fumigatus* secretome are secreted via classical pathway and their GO analysis revealed involvement in carbohydrate metabolic processes and regulation of host immune response, while the remaining 8 are secreted via a non-classical pathway and their GO analysis revealed involvement in response to mycotoxin, synthesis of mycotoxin and gliotoxin, and response to oxidative stress apart from carbohydrate metabolic processes. Within the cell membrane proteins of *A. fumigatus,* 151 out of 1129 proteins (13.3%) were upregulated during pathogenesis, and GO analysis revealed that many of them are involved in transmembrane transport processes. Moreover, 45 SSPs in the *A. fumigatus* secretome, of which 21 have effector-like features, were upregulated over 2-fold during pathogenesis. For example, Afu3g14940 a predicted peptidase inhibitor I78 family protein which has a known conserved effector motif ([KRHQSA][DENQ]EL) and Afu1g01210 with the adhesion fasciclin domain, are among the 21 effector-like SSPs with over 2-fold upregulation during pathogenesis.

### Conservation of secretome across *Aspergillus* species

The ability of multiple *Aspergilli* to switch into an opportunistic pathogen may be derived from conserved proteins. Conversely, species-specific proteins may explain the observed differences in virulence between *Aspergilli.* To determine the unique and conserved proteins across *Aspergillus* species, OrthoMCL^84^ was used to identify orthologs between the proteins of nine *Aspergillus* species. As *A. ustus* has an incomplete genome, it was omitted from this comparative analysis. Figures 4A-C display for each species the fraction of unique and conserved proteins within the complete proteome, cell membrane proteins and secretomes, respectively, across the nine *Aspergillus* species. In comparison to the cell membrane proteins, the secretomes of *Aspergillus* species have a larger set of unique proteins and a smaller set of conserved proteins across the nine *Aspergillus* species. Interestingly, 3.6% of the secretome of *A. fumigatus* is unique, while 27.3% and 26.9%, respectively, of the secretomes of *A. niger* and *A. glaucus* is unique, when compared across the nine *Aspergillus* species. Thus, the secretome of *A. fumigatus* has among the smallest fraction of unique proteins which is in contrast to its position as the most prevalent agent of aspergillosis. The unique and conserved proteins identified here across the secretomes of *Aspergillus* species could lead to a better understanding of the important players for virulence and provide potential targets for the development of broad spectrum antifungals.

**Figure 4:**
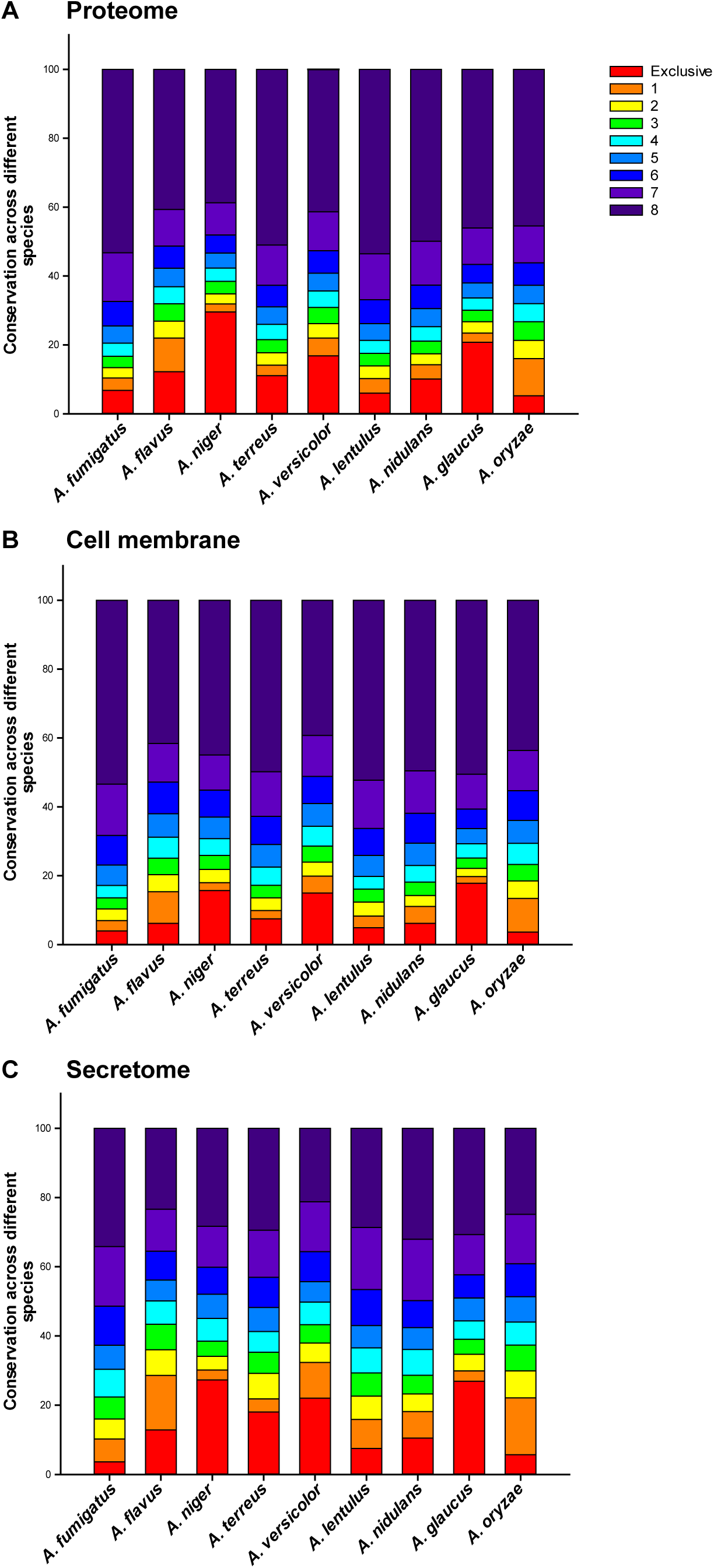
Conservation of **(a)** complete proteome, **(b)** the set of cell membrane proteins, and **(c)** the set of secreted extracellular proteins across the nine *Aspergillus* species considered here. The fraction of proteins that are unique to a specific *Aspergillus* species is shaded in orange at the bottom of the stacked bar chart while the fraction of proteins conserved across all nine *Aspergillus* species are shaded in dark violet at the top of the stacked bar chart.

### Comparison with the human proteome

Ideally, antifungals should target fungi-specific proteins to prevent cross-toxicity with human proteins. In order to identify fungi-specific druggable targets, BLASTP with an E-value cut-off of ≤ 1e-3 was used to identify *Aspergillus* proteins without a close homolog in the human proteome. We find that more than 70% of the secretomes in *Aspergillus* species is unique, without a homolog in humans (Figure 5). In contrast to the secretomes, the cell membrane fraction or whole proteome of *Aspergillus* species have a much lower fraction of unique proteins without a homolog in humans (Figure 5). Specifically, *A. fumigatus* has 71% of its secretome and 52% of its cell membrane proteins without a homolog to humans, and this offers a significant candidate set that may be screened for druggability or biomarker design. The striking difference between conservation of whole proteome and secretome of *Aspergillus* species in humans can likely be attributed to the large presence of proteins associated with plant cell wall degradation^85^ in the *Aspergillus* secretomes.

**Figure 5:**
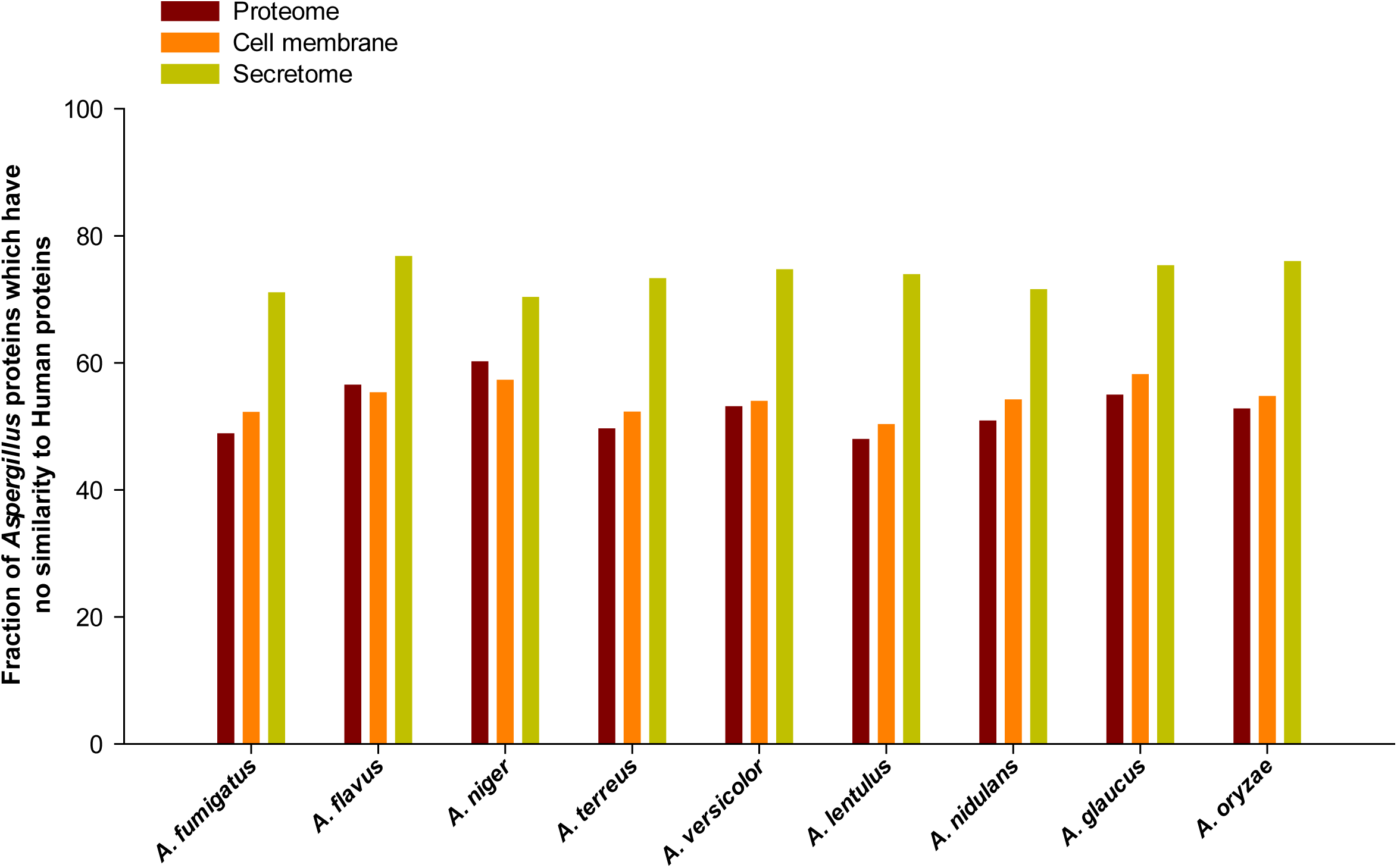
Fraction of the complete proteome, set of cell membrane proteins and set of secreted extracellular proteins in *Aspergillus* species without a close sequence homolog in human proteome.

### Antigenicity of the secretome

Protein vaccines have been proven successful for several invasive fungi^86,87,88^. For example, vaccination with the *A. fumigatus* allergen Aspf3 protected immunosuppressed mice from developing aspergillosis^89^. One measure of clinical importance is the number of potential antigenic regions on a protein. The more antigenic a protein the more likely it can be used as a biomarker, targeted for immunotherapy, or used in vaccinations. To help prioritize *Aspergillus* proteins, the Abundance of Antigenic Regions (AAR)^24^ value of proteins was calculated using two methods, Kolaskar-Tongaonkar^90^ and BepiPred 2.0^91^ (Methods). The lower the AAR value the more antigenic the protein^24^. Interestingly, in each *Aspergillus* species, the average AAR of the secretome is always significantly lower than cell membrane proteins (Figure 6; Methods). Furthermore, the average AAR of the secretome and cell membrane proteins was also found to be significantly lower (p < 0.001) and significantly higher (p < 0.001), respectively, in comparison to the average AAR of randomly chosen, equally sized sets of proteins from the whole proteome (Methods). Note that similar observations on relative AAR of secretome and cell membrane proteins were also recently made in tapeworms, *Taenia solium*^24^ and *Echinococcus multilocularis*^92^, and in bacterium *Mycobacterium tuberculosis*^93^. Thus, secreted proteins are likely to be more antigenic than cell membrane proteins, and while choosing candidates for vaccine development, the secretome may provide a more antigenic landscape than cell membrane proteins. While a comparison of the AAR of secretome and cell membrane proteins in *Aspergillus* species provides insight on relative virulence within a proteome, no significant difference in the average AAR could be detected between *A. fumigatus* and the *Aspergillus* cohort.

**Figure 6:**
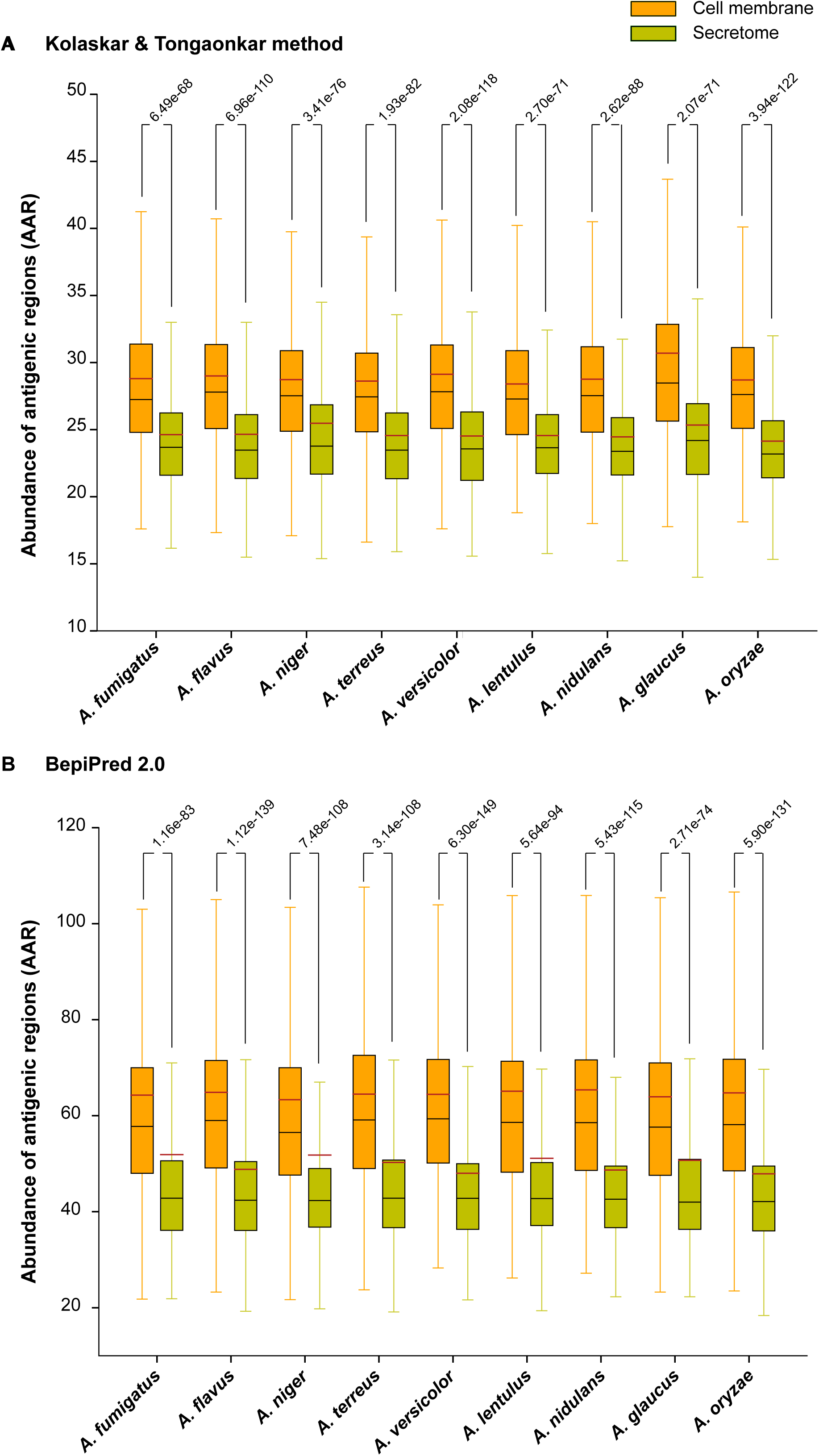
Distribution of Abundance of Antigenic Regions (AAR) values for the set of cell membrane proteins and secreted extracellular proteins in *Aspergillus* species considered here. AAR values were computed using two different methods: **(a)** Kolaskar-Tongaonkar^90^ method and **(b)** BepiPred 2.0^91^. In each box plot, the lower end of the box represents the first quartile, black line inside the box is the median, brown line is the mean and the upper end of the box represents the third quartile of the distribution. In this figure, we also report the p-value from the comparison of the distributions of AAR values in the set of cell membrane proteins and in the set of secreted proteins for each *Aspergillus* species performed using Wilcoxon rank-sum test.

### Druggability analysis of *A. fumigatus* secretome and cell membrane proteins upregulated during pathogenesis

Beyond immune-therapy, biomarker, and vaccine development, drugs are often the main line of defence against aspergillosis. However, the current arsenal to combat aspergillosis is limited to 7 approved drugs. Furthermore, resistance to these few drugs is an ever-emerging threat^14^. Particularly, fungi are adept at developing resistance to drugs that act on intracellular proteins by pumping them back out into the extracellular matrix through efflux pumps. However, targeting of secreted proteins and cell membrane proteins could circumvent this specific defence mechanism altogether. To expedite drug discovery pipeline, DrugBank^25^ provides a database of known drugs and their targets, many of which have been successfully repurposed against similar protein targets in different pathogens^94^.

Firstly, the list of 4063 known drug target proteins was compiled from DrugBank^25^. Secondly, the secreted and cell membrane proteins in *Aspergillus* species with no close human homologs based on BLASTP with an E-value cut-off of ≤ 1e-3 were determined. Thereafter, secreted and cell membrane proteins in *Aspergillus* species with no close human homologs but with sequence similarity to known drug target proteins based on BLASTP E-value cut-off of ≤ 1e-5 were determined (Supplementary Table S5). In *A. fumigatus,* 50 secreted and 16 cell membrane proteins were found to have sequence similarity with known drug target proteins. Subsequently, we focused on upregulated genes in a transcriptome dataset^26^ for *A. fumigatus* during pathogenesis to identify potential target proteins for drug repurposing.

In the transcriptome dataset^26^ for *A. fumigatus*, we found 97 secreted and 101 cell membrane proteins with no close human homologs to be significantly upregulated over 2-fold during pathogenesis. These 97 secreted and 101 cell membrane proteins in *A. fumigatus* were searched against known target protein sequences for sequence similarity to known drug targets. Seven secreted proteins, Afu7g06140, Afu8g01670, Afu2g15160, Afu5g14380, Afu6g09740, Afu3g00610 and Afu1g09900, of which three proteins, Afu8g01670, Afu6g09740 and Afu1g09900, are secreted via non-classical pathways, and one membrane protein, Afu6g03570, had a significant hit to known drug target proteins. After establishing the sequence similarity between *A. fumigatus* proteins and known drug target proteins, structural similarity was probed (Methods). Experimentally identified protein structure was available only for Afu6g09740, and it was downloaded from the protein data bank^95^ (PDB). As good-quality structure model was also unavailable for Afu2g15160, it was omitted from later analysis. For the remaining five secreted proteins and one cell membrane protein, structure models were obtained from ModBase^96^ and SWISS-MODEL^97^. Thereafter, the compiled protein structures were compared to their DrugBank counterparts for structural similarity. Four secreted proteins, Afu5g14380, Afu8g01670, Afu1g09900 and Afu6g09740, had significant structural similarity with TM scores greater than 0. 8 and root-mean-square deviation (RMSD) values lower than 3Å^2^ (Table 1; Methods). One of the proteins, Afu8g01670, is a putative bifunctional catalase-peroxidase which has been shown to be involved in virulence^98^. Afu5g14380 is an α-glucuronidase involved in the hydrolysis of xylan. Afu1g09900 is involved in degradation of arabinoxylan. Afu6g09740 is a thioredoxin reductase which is a part of the gene cluster involved in biosynthesis of gliotoxin in *A. fumigatus,* and its knockout has been shown to affect the oxidation of gliotoxin and render the fungi hypersensitive to exogenous gliotoxin^99^. Afu6g09740 was also found to be immunogenic in humans and a potential biomarker of aspergillosis^100^.

**Table 1:** Evaluation of the structural similarity of secreted and cell membrane proteins upregulated in the transcriptome of *Aspergillus fumigatus* Af293 with their close sequence homologs among known drug target proteins in DrugBank^25^. Note that the *A. fumigatus* proteins shaded in gray are not structurally similar to their close sequence homologs among known drug target proteins.

| Protein identifier | Log<sub>2</sub> fold change in transcriptome | Drug target protein identifier | TM score | RMSD (Å<sup>2</sup>) |
| --- | --- | --- | --- | --- |
| Afu5g14380 | 1.4 | Q8VVD2 | 0.98 | 0.19 |
| Afu8g01670 | 2.3 | Q939D2 | 0.99 | 0.28 |
|  |  | P9WIE5 | 0.97 | 0.72 |
| Afu1g09900 | 1.1 | Q9XBQ3 | 0.91 | 0.99 |
| Afu6g09740 | 1.3 | P66010 | 0.86 | 2.68 |
| Afu7g06140 | 7.2 | P40406 | 0.73 | 4.05 |
| Afu6g03570 | 1.3 | P40406 | 0.72 | 3.29 |
| Afu3g00610 | 1.2 | P05618 | 0.26 | 15.24 |
|  |  | P26827 | 0.25 | 19.25 |

Given the sequence and structural similarity of these four *A. fumigatus* proteins, Afu5g14380, Afu8g01670, Afu1g09900 and Afu6g09740, to known drug targets, the proteins were subsequently tested whether they could bind to their respective drugs (Table 2; Methods). The four proteins were analyzed for the presence of ligand binding pockets using metaPocket 2.0^101^. For each of the four proteins, the top three ligand binding pockets were tested for ligand binding affinity using AutoDock Vina^102^. Table 2 lists the four *Aspergillus* proteins, the corresponding target proteins in DrugBank with both sequence and structural similarity, their reported approved or experimental drugs, and the binding affinity of each drug to the ligand pockets. Using AutoDock Vina^102^, it was found that most of the identified drug candidates have an affinity value of ≤ -5.0 kcal/mol for their respective *Aspergillus* proteins which signifies good binding and suggests that the drugs may be able to bind to the upregulated and secreted *A. fumigatus* proteins (Table 2; Methods). Further experimental validation of these hit molecules is needed to verify whether the drug will act upon the *A. fumigatus* protein in the same fashion and whether the drug will have an impact upon the ability of *A. fumigatus* to cause aspergillosis.

**Table 2:** Binding affinity of potential drugs against predicted pockets of the four candidate proteins in *Aspergillus fumigatus.* Note that the drug candidates were identified based on sequence and structural similarity of the four upregulated proteins in *A. fumigatus* with known drug target proteins in DrugBank^25^ database. The ligand binding pockets were predicted using metaPocket 2.0^101^ webserver and binding affinities of drugs to proteins were computed using AutoDock Vina^102^ (Methods).

| Protein identifier | Protein identifier of drug target protein | Drug identifier in DrugBank | Drug name | Drug class | Mean binding affinity (kcal/mol) |  |  |
| --- | --- | --- | --- | --- | --- | --- | --- |
|  |  |  |  |  | First pocket | Second pocket | Third pocket |
| Afu8g01670 | P9WIE5 | DB00609 | Ethionamide | Approved | -5.9 ± 0.0 | -5.9 ± 0.0 | -5 ± 0.0 |
|  |  | DB00951 | Isoniazid | Approved | -5.53 ± 0.015 | -5.91 ± 0.01 | -5.3 ± 0.0 |
|  | Q939D2 | DB08638 | 2-Amino-3-(1-hydroperoxy-1H-indol-3-yl)propan-1-ol | Experimental | -7.6 ± 0.033 | -7.6 ± 0.0 | -5.87 ± 0.015 |
| Afu5g14380 | Q8VVD2 | DB03156 | D-Glucuronic Acid | Experimental | -5.1 ± 0.0 | -6.5 ± 0.0 | -5.6 ± 0.0 |
|  |  | DB04303 | 4-O-methyl-alpha-D-glucuronic acid | Experimental | -5 ± 0.0 | -6.3 ± 0.0 | -5.3 ± 0.0 |
|  |  | DB02722 | 4-O-methyl-beta-D-glucuronic acid | Experimental | -5 ± 0.0 | -5.8 ± 0.0 | -5.11 ± 0.01 |
| Afu6g09740 | P66010 | DB00548 | Azelaic Acid | Approved | -5.53 ± 0.026 | -5.52 ± 0.065 | -5.21 ± 0.031 |
| Afu1g09900 | Q9XBQ3 | DB03196 | 4-Nitrophenyl-Ara | Experimental | -6.3 ± 0.0 | -6 ± 0.0 | -5.54 ± 0.031 |
|  |  | DB03870 | Ara-Alpha(1,3)-Xyl | Experimental | -5.51 ± 0.023 | -5.54 ± 0.016 | -5.62 ± 0.029 |

### Comparison of our prediction pipeline with earlier work

We have designed here a computational prediction pipeline to identify the secretome and cell membrane proteins in *Aspergillus* species, and the pipeline can be used for any fungi. To our knowledge, FSD^16^ and FunSecKB2^17^ are the only databases on pan-fungi secretome prediction. While FSD and FunSecKB2 contain pre-computed secretome predictions for several fungal genomes, SECRETOOL^18^ provides an online sever that enables implementation of a computational pipeline to predict secreted proteins among user-input protein sequences. In Supplementary Text and Supplementary Table S6, we report a detailed comparison of the secretome predictions from our pipeline with those from FSD, FunSecKB2 and SECRETOOL.

Based on this comparative analysis with FSD, FunSecKB2 and SECRETOOL, our pipeline has following additional features. Firstly, we incorporate available experimental information from high-throughput studies and UniProt on secreted proteins (Branch A in Figure 1). Secondly, we incorporate and prioritize UniProt annotations with experimental evidence for presence of signal peptide, GPI anchor or TM domain in a protein sequence over bioinformatic prediction tools (Branch B in Figure 1). Thirdly, we provide a sub-classification of the secreted extracellular proteins into those sorted by classical pathway and those transported via non-classical pathways. Fourthly, we have used an alternate approach whereby orthologs to secreted proteins with experimental evidence in other fungi is used to predict secretion via non-classical pathways (Branch C in Figure 1). We remark that our pipeline predicts a much smaller set of secreted proteins via non-classical pathways (albeit with much higher confidence) in comparison to FSD which uses SecretomeP^74,75^, a tool designed for bacteria and mammals rather than fungi. Lastly, to our knowledge, this is the first study to perform a comparative analysis of both secreted and cell membrane proteins across *Aspergillus* species causing aspergillosis.

## Conclusions

Aspergillosis is a serious concern among immune-compromised patients worldwide. Fungal secretome and cell membrane proteins are often the first point of contact between fungal-host interactions, and thus are key factors in initiating infection or to mounting any defence against the disease. Thus, here, we report a comprehensive set of secreted and cell membrane proteins in *Aspergillus fumigatus* and nine other *Aspergillus* species that will serve as a valuable resource for understanding the pathogenesis of aspergillosis. We also identified protein cohorts that could be further screened and targeted for development of novel treatments. To summarize, we have firstly implemented a computational pipeline that integrates data from experiments and bioinformatic-based predictions to identify a comprehensive set of secreted and cell membrane proteins. Secondly, the identified secretome was analyzed for SSPs and effector-like SSPs for correlating their role during pathogenesis. Thirdly, the antigenicity of the secretome and cell membrane proteins were computed to aid in understanding their role in inducing immune responses. Fourthly, the secreted and cell membrane proteins without homologs in humans were scanned for similarity with known drug target proteins in DrugBank to identify new druggable proteins. Lastly, a systematic representative drug repurposing analysis was performed in *A. fumigatus* responsible for overwhelming majority of the aspergillosis cases. Contextualization of an expression dataset for *A. fumigatus* within its secretome and cell membrane proteins led to identification of potential targets with over 2-fold upregulation during pathogenesis, and similar to DrugBank target proteins based on both sequence-wise and structure-wise comparison, and able to dock with their respective drug. This led to identification of new potential drug candidates against aspergillosis from existing drugs. The identified secretome and cell membrane proteins in the ten *Aspergillus* species along with their sub-classification and functional annotations is also hosted at https://cb.imsc.res.in/aspertome for convenient access and retrieval.

## Methods

### Protein sequences and associated annotations for *Aspergillus* species

The proteomes of *A. fumigatus* Af293, *A. niger* CBS 513.88, *A. terreus* NIH2624, *A. versicolor* CBS 583.65, *A. lentulus* IFM 54703T, *A. nidulans* FGSC A4, *A. glaucus* CBS 516.65 and *A. oryzae* RIB40 were retrieved from the *Aspergillus* genome database (AspGD^103^; http://www.aspgd.org/). The proteomes of *A. flavus* NRRL3357 and *A. ustus* 3.3904 were retrieved from FungiDB^104^ (http://fungidb.org/fungidb/) and Ensembl Genomes^105^ (http://ensemblgenomes.org/), respectively. Note that the *A. ustus* 3.3904 genome sequence is still incomplete, and thus, its incomplete proteome was used here for predictions.

### Experimentally identified secreted proteins

We performed an extensive literature search to compile 46 published high-throughput experimental studies^27-73^ which have characterized the secretome of 6 *Aspergillus* species studied here (Supplementary Table S1). Note that protein identifiers in these experimental studies for the reference strain or other strains of the same *Aspergillus* species were mapped to standard identifiers in the reference sequence using OrthoMCL^84^ (http://orthomcl.org/orthomcl/) with inflation option set at 1.5. In addition to high-throughput studies, experimentally verified secreted or cell membrane proteins in *Aspergillus* species were retrieved from UniProt^19^ (http://www.uniprot.org/) by filtering proteins with subcellular localization as extracellular or cell membrane, respectively, with evidence code of ECO:0000269. Note that ECO:0000269 corresponds to manually curated information with published experimental evidence. Similarly, experimentally verified secreted proteins in all other fungal species were retrieved from UniProt^19^ by filtering proteins with subcellular localization as extracellular with evidence code of ECO:0000269. OrthoMCL^84^ with inflation option set at 1.5 was used to identify orthologs of experimentally verified secreted proteins in other fungi in *Aspergillus* species.

### Bioinformatic-based predictions

The following computational tools were employed to predict protein-based features in our secretome prediction pipeline. The presence of a signal peptide in the N-terminus was predicted using SignalP 4.1^106^ (http://www.cbs.dtu.dk/services/SignalP/) and Phobius^107^ (http://phobius.sbc.su.se/). The presence of GPI anchor was predicted using PredGPI^108^ (http://gpcr.biocomp.unibo.it/predgpi/) and big-PI^109^ (http://mendel.imp.ac.at/gpi/fungi_server.html). The presence of TM domain was predicted using TMHMM 2.0^110^ (http://www.cbs.dtu.dk/services/TMHMM/) and Phobius^107^. Note that TM domain predictions by TMHMM within the last 70 amino acid residues of the N-terminus of a protein sequence were not considered as the tool can sometimes predict signal peptides as false-positive TM domains. The presence of ER retention signal was predicted using PS SCAN^111^ with PROSITE (https://prosite.expasy.org/) pattern PS00014^112^. The subcellular localization of proteins was predicted using WoLF PSORT 0.2^113^ (https://psort.hgc.jp/), TargetP 1.1^114^ (http://www.cbs.dtu.dk/services/TargetP/) and ProtComp 9^115^ (http://www.softberry.com/berry.phtml?topic=protcomp&group=help&subgroup=proloc). Importantly, UniProt^19^ annotations with published experimental evidence (ECO:0000269) for the protein-based features were also retrieved for *Aspergillus* proteins. UniProt identifiers for *Aspergillus* proteins were mapped to reference sequence identifiers using pre-existing maps from AspGD^103^, UniProt^19^, and in-house python scripts.

While combining information from different sources to decide on the presence of signal peptide or GPI anchor or TM domain, a decision is made based on tool predictions using an inclusive rule if UniProt annotation is not available, else decision is made only on UniProt annotation by overriding tool predictions. While combining information from different sources to decide on subcellular localization, the decision is made based on tool predictions using a majority rule if UniProt annotation is not available, else decision is made only on UniProt annotation by overriding tool predictions. Thus, our pipeline gives precedence to published experimental evidence over bioinformatic-based predictions. Supplementary Table S2 contains tool-based predictions and UniProt annotations for secreted and cell membrane proteins in *Aspergillus* species.

### Functional annotation of secreted proteins

The predicted secretomes in *Aspergillus* species were annotated with information on protein family from Pfam^116^ database (http://pfam.xfam.org/) and carbohydrate-binding modules from CAZy^117^ (http://www.cazy.org/)^117^ and dbCAN v5.0^118^ (http://csbl.bmb.uga.edu/dbCAN/) databases using hmmpress and hmmscan utilities in HMMER3 (http://hmmer.org/) with profile-specific GA (gathering) thresholds. Furthermore, the predicted secretomes in *Aspergillus* species were annotated with protein family and domain information from TIGRFAM^119^ (http://www.jcvi.org/cgi-bin/tigrfams/index.cgi), SFLD^120^ (http://sfld.rbvi.ucsf.edu/django/), SMART^121^ (http://smart.embl-heidelberg.de/), CDD^122^(https://www.ncbi.nlm.nih.gov/cdd), PROSITE^112^, SUPERFAMILY^123^ (http://supfam.org/SUPERFAMILY/), PRINTS^124^ (http://130.88.97.239/PRINTS/index.php), PANTHER^125^ (http://www.pantherdb.org/), COILS^126^ (https://embnet.vital-it.ch/software/COILS_form.html) and MobiDB-lite^127^ (http://protein.bio.unipd.it/mobidblite/) accessed through InterPro version 64.0 (https://www.ebi.ac.uk/interpro/) using InterProScan 5^128,129^. In addition, the secreted proteins in *Aspergillus* species were annotated with GO terms using FungiFun2^130^ (https://elbe.hki-jena.de/fungifun/fungifun.php). Supplementary Table S3 contains detailed annotations for secreted and cell membrane proteins in *Aspergillus* species.

### Identification of small secreted and effector-like proteins

In the predicted secretomes of *Aspergillus* species, SSPs were defined as those with sequence length less than or equal to 300 amino acid residues^21,82,131^. The SSPs were then evaluated for effector-like properties based on their EffectorP^81^ (http://effectorp.csiro.au/) predictions or cysteine-richness. For identifying effector-like SSPs, cysteine-rich sequences are those which contain at least 4 cysteine residues and have greater than 5% of their total amino acid residues as cysteines^20,132^. The predicted SSPs in the secretomes of *Aspergillus* species were also scanned for known fungal or oomycetes effector motifs^80^ (DEER, RXLR, RXLX[EDQ], [KRHQSA][DENQ]EL, [YW]XC and RSIVEQD) using the FIMO package in MEME program suite^133^ with E-value cut-off of ≤ 1e-4 (Supplementary Table S4).

### Antigenicity of secreted proteins

Antigenic regions in *Aspergillus* proteins were predicted using Kolaskar-Tongaonkar^90^ method implemented in EMBOSS package^134^ with a threshold of 1.0 and BepiPred 2.0^91^ (http://www.cbs.dtu.dk/services/BepiPred/index.php) with the default threshold of 0.5. Note that only predicted antigenic regions with a length ≥ 6 amino acids were accounted in the later analysis. For each protein in *Aspergillus* species, the AAR value was computed following the method proposed by Gomez *et al*^24^. The AAR value of a given protein is computed by dividing the length of its amino acid sequence by the number of predicted antigenic regions. The average AAR value was computed for the set of secreted and cell membrane proteins, respectively, in each of the *Aspergillus* species considered here (Figure 6). To test whether the computed average AAR values for the set of secreted and cell membrane proteins, respectively, were significantly different from the average AAR value for the whole proteome of the same species, a p-value was computed by comparing against the average AAR values for 1000 equally-sized sets of randomly drawn proteins from the complete proteome of the organism. To test whether the average AAR value for the set of secreted proteins is significantly different from that of the set of cell membrane proteins of an *Aspergillus* species, Wilcoxon rank-sum test was performed to compare the two sets of different sizes (Figure 6).

### Identification of candidate drug targets in secreted proteins

A microarray dataset for *A. fumigatus* Af293 from infected murine lungs^26^ was obtained from Array Express (Accession number E-TABM-327). The microarray dataset from infected murine lungs^26^ gave a reliable signal for 9121 genes in *A. fumigatus* Af293 which covers more than 90% of the secretome and 88% of the cell membrane proteins in the fungus. Gene expression data^26^ for *A. fumigatus* Af293 was next analyzed within the context of the predicted secreted and cell membrane proteins to identify differentially expressed proteins with over 2-fold upregulation in each fraction during pathogenesis for further analysis. Thereafter, secreted and cell membrane proteins in *A. fumigatus* Af293 with over 2-fold upregulation during pathogenesis and sequence similarity with known drug target proteins from DrugBank^25^ database were identified using BLASTP (https://blast.ncbi.nlm.nih.gov/Blast.cgi) with E-value cut-off of ≤ 1e-5 (as specified by DrugBank database).

To further ascertain the similarity between the filtered set of *A. fumigatus* proteins in secretome and cell membrane fraction with over 2-fold upregulation during pathogenesis and with sequence similarity with known drug target proteins from DrugBank^25^, the structural similarity between filtered proteins and their sequence-based homologs among known drug targets was assessed. Structure of proteins from x-ray diffraction experiments were retrieved from PDB^95^ (https://www.rcsb.org/pdb/home/home.do). However, experimentally identified protein structures are not available for most of the filtered set of secreted and cell membrane proteins in *A. fumigatus* Af293, and in such cases, the protein structures modelled using close homologs (>40% sequence identity) from ModBase^96^ (https://modbase.compbio.ucsf.edu/modbase-cgi/index.cgi) and SWISS-MODEL^97^ (https://swissmodel.expasy.org/) were used to evaluate structural similarity. The modelled structures of filtered proteins in *A. fumigatus* Af293 were compared to the structure of their corresponding sequence-based homolog among the known drug target proteins in DrugBank^25^ based on TM score computed using TM Align^135^ (https://zhanglab.ccmb.med.umich.edu/TM-align/) and root-mean-square deviation (RMSD) value between locally aligned atoms computed using PyMOL (The PyMOL Molecular Graphics System, Version 1.8 Schrödinger, LLC). Filtered proteins in *A. fumigatus* Af293 with significant structural similarity to known drug target proteins were subsequently tested for binding by the respective drug.

The ligand binding pockets in the protein structure model for the filtered proteins in *A. fumigatus* Af293 were predicted using metaPocket 2.0^101^ (http://projects.biotec.tu-dresden.de/metapocket/). For docking drugs to the predicted protein pockets, both protein and drug molecule were prepared by adding explicit hydrogen atoms and cleaned up using python scripts^136^, prepare_receptor4.py and prepare_ligand4.py, with the default option. AutoDock Vina^102^ (http://vina.scripps.edu/) was used for docking the drug against the predicted pockets in proteins with the option for exhaustiveness set at 24 to obtain consistent results^137^. Lastly, the binding affinity results obtained by docking drugs to the protein pockets from AutoDock Vina^102^ were tabulated (Table 2).

## Data availability

All data generated or analyzed during this study is included in this article and its supplementary information.

## Acknowledgements

We thank Aashish Shivkumar for help in data collection, and B. Raveendra Reddy and P. Mangalapandi for help with webserver for hosting our dataset. AS would like to acknowledge financial support from the Department of Science and Technology (DST) India through the award of a start-up grant (YSS/2015/000060) and Ramanujan fellowship (SB/S2/RJN-006/2014), and Max Planck Society Germany through the award of a Max Planck Partner Group on Mathematical Biology.

## Author contributions

AS conceived the project. AS, RPV, JPC designed research including the computational prediction pipeline. RPV, KM, MV compiled and curated data from various sources. RPV implemented the computational prediction pipeline. RPV, AJ, JPC, AS analyzed data. RPV, JPC, AS wrote the manuscript. All authors have read and approved the manuscript.

## Conflicts of interest

The authors declare that they have no conflicts of interest.

## Supplementary Information

**Supplementary Table S1:** Complied list of experimentally determined secreted proteins in different *Aspergillus* species from high-throughput proteomic studies.

**Supplementary Table S2:** Experimentally identified and computationally predicted secreted extracellular proteins and cell membrane proteins in different *Aspergillus* species. In this table, we have also included information on UniProt annotations with published experimental evidence, predictions from different computational tools, and Abundance of Antigenic Regions (AAR) score for each protein.

**Supplementary Table S3:** Functional annotation of identified secreted proteins and cell membrane proteins in different *Aspergillus* species.

**Supplementary Table S4:** Small secreted proteins (SSPs) among the secretomes of different *Aspergillus* species.

**Supplementary Table S5:** List of secreted and cell membrane proteins in *Aspergillus* species without sequence similarity to human proteins but with sequence similarity to known drug targets in other organisms. In this table, we have also compiled the drugs associated with each target protein from the DrugBank database. In the first sheet, we list the number of secreted and cell membrane proteins across *Aspergillus* species which have sequence similarity to known drug targets from DrugBank database.

**Supplementary Table S6:** Comparison of secretome predictions from our computational pipeline in *Aspergillus* species with predictions from earlier work. In the first sheet, we list the subset of *Aspergillus* species considered here for which FSD and FunSecKB2 have predictions in their database. In the second sheet, we compare the bioinformatic tools employed in our prediction pipeline with those in FSD, FunSecKB2 and SECRETOOL. In the third sheet, we compare predictions from our pipeline with those from FSD. In the fourth sheet, we compare predictions from our pipeline with those from FunSecKB2. In the fifth sheet, we compare predictions from our pipeline with those from SECRETOOL.

